# Examining oculomotor behavior in central vision loss with a gaze-contingent display

**DOI:** 10.1101/2025.10.12.681883

**Authors:** Marcello Maniglia, Jason Vice, Elliot Maxwell, Kristina M Visscher, Aaron R Seitz

## Abstract

Patients with central vision loss due to macular degeneration (MD) must rely on their peripheral vision for tasks normally performed by the fovea. Many patients develop a preferred retinal locus (PRL), an eccentric retinal location used as a substitute for the damaged fovea in tasks such as face recognition, navigation, and reading. However, the mechanisms underlying PRL development remain elusive, and no single hypothesis fully explains its characteristics. Investigations into PRL development are hindered by oculomotor assessments, which often focus on fixation ability while neglecting other eye movement characteristics and potentially conflating different behaviors over time. In previous work, we introduced a series of oculomotor metrics in cases of simulated central vision loss, demonstrating that complex profiles of eye movement behavior can be extracted from a simple visual task. Here we present longitudinal data from 10 patients with MD as evidence of the feasibility of using these metrics to characterize different profiles of eye movements following central vision loss. Consistent with findings in healthy individuals using artificial scotoma, the metrics reveal substantial individual differences in behavior, both at baseline and after visual training. Overall, patients exhibit significantly higher saccadic re-referencing than controls, despite larger inter-individual differences. These metrics provide a detailed evaluation of oculomotor behavior in patients with central vision loss and offer a valuable tool for assessing progress in training protocols.

## Introduction

Macular degeneration (MD), a prevalent retinal condition among older adults, is the leading cause of visual impairment in the Western world ^1^. The public health impact of MD is exacerbated by the fact that the most common form is age-related macular degeneration, which is coupled with the aging population globally. Early stages of MD are characterized by retinal distortions in the center of the visual field, which often go unnoticed until the pathology reaches advanced stages. Advanced stages usually result in a central retinal scotoma leading to a complete loss of central vision. Currently, there is no standard intervention for MD, and rehabilitation efforts are complicated by the heterogeneous nature of the condition. Variations in lesion size, shape and location, comorbidity, and demographic characteristics further add to the challenge.

Understanding eye movement behavior in MD is paramount to characterizing compensatory oculomotor strategies, evaluating spontaneous adaptation, and assessing the effectiveness of rehabilitation approaches. In this study, we present a novel approach to eye movement analysis under conditions of central vision loss. This approach aims to disentangle differences in behaviors that might be conflated in standard oculomotor analyses.

### PRL Development and Challenges in Analysis

Patients with MD often adopt compensatory oculomotor strategies by developing a Preferred Retinal Locus (PRL), a peripheral retinal region that patients use to perform tasks otherwise reliant on the fovea ^2–5^. This surrogate fovea has been observed in approximately 70%-80% of patients with MD ^6,7^. Growing evidence suggests that characteristics of the PRL might be influenced by multiple factors, including the task and stimuli used during eye exams. Some studies suggest that a PRL might not necessarily imply a shift in oculomotor reference (e.g., saccadic eye movements may not be directed to that new fixation target). Furthermore, different tests, such as static versus dynamic fixation tasks, might reveal different types of oculomotor referencing ^8^. Specifically, Chung (2013) distinguishes between ‘fixational’ saccades and ‘re-fixational’ saccades, where the first are automatic and the second volitional, and argues that patients might exhibit re-referencing for one type of eye movement but not the other.

Further, evidence suggests that certain patients with MD may exhibit multiple PRLs ^2,9–13^. As a result, the most common PRL analyses, which typically presume a single PRL with a unimodal distribution of fixation data, risk conflating diverse PRLs into a singular distribution ^14^. Consequently, the derived PRL estimate, typically assessed via a bivariate contour ellipse area (BCEA), could be misleading both in indicating the level of fixation stability, as well as in providing the exact retinal location of the PRL. Traditional low vision eye exams rely on expensive and specialized equipment, such as scanning laser ophthalmoscopy (SLO), optical coherence tomography (OCT) and microperimetry, which limits widespread application, posing feasibility challenges to our ability to collect large samples of patient data.

### Oculomotor Metrics for MD Analysis

Challenges related to task design, stimulus selection, data analysis approaches, and access to specialized equipment hinder our understanding of compensatory oculomotor behavior in central vision loss, including characteristics of the PRL. Recent technological advancements, such as high-resolution, video-based eye trackers can provide more accessible equipment for the evaluation of eye movement behavior in low vision, enabling the use of a wider range of tasks and stimuli to assess the PRL. We recently proposed a series of oculomotor metrics to better classify eye movements after central vision loss beyond the standard fixation analysis (details provided in Maniglia et al. (2020) and summarized below and in Figure 1). This includes additional metrics, such as temporal connotations of eye movements and ‘useful’ trials in a multiple-trial assessment ^15^. Specifically, we proposed six metrics to characterize compensatory eye movements to be collected over a multi-trial test:

**Figure 1.**
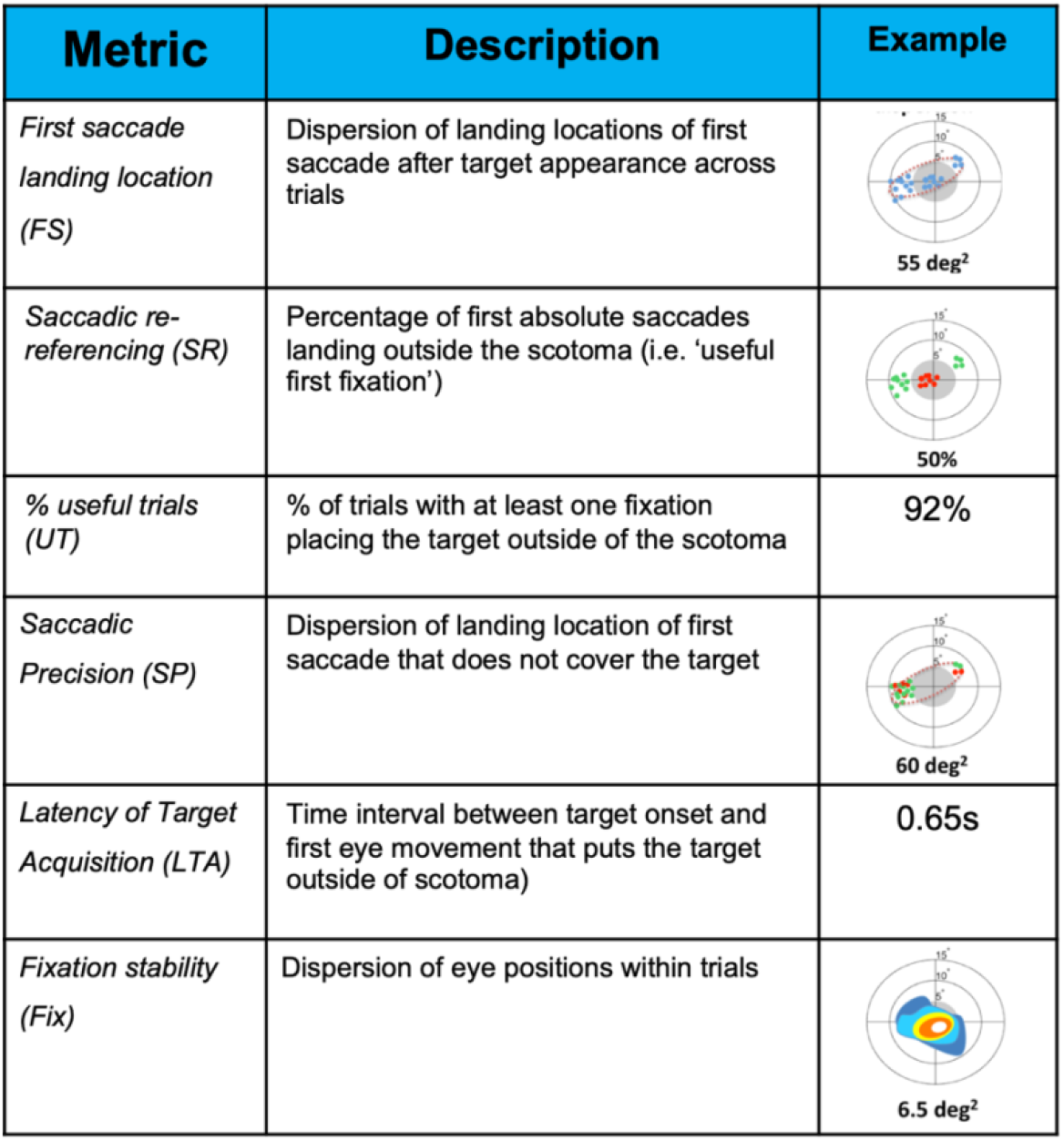
Summary description of the oculomotor metrics used in this study to evaluate oculomotor behavior in conditions of central vision loss (from Maniglia, Visscher and Seitz, 2020)

**Figure 2.**
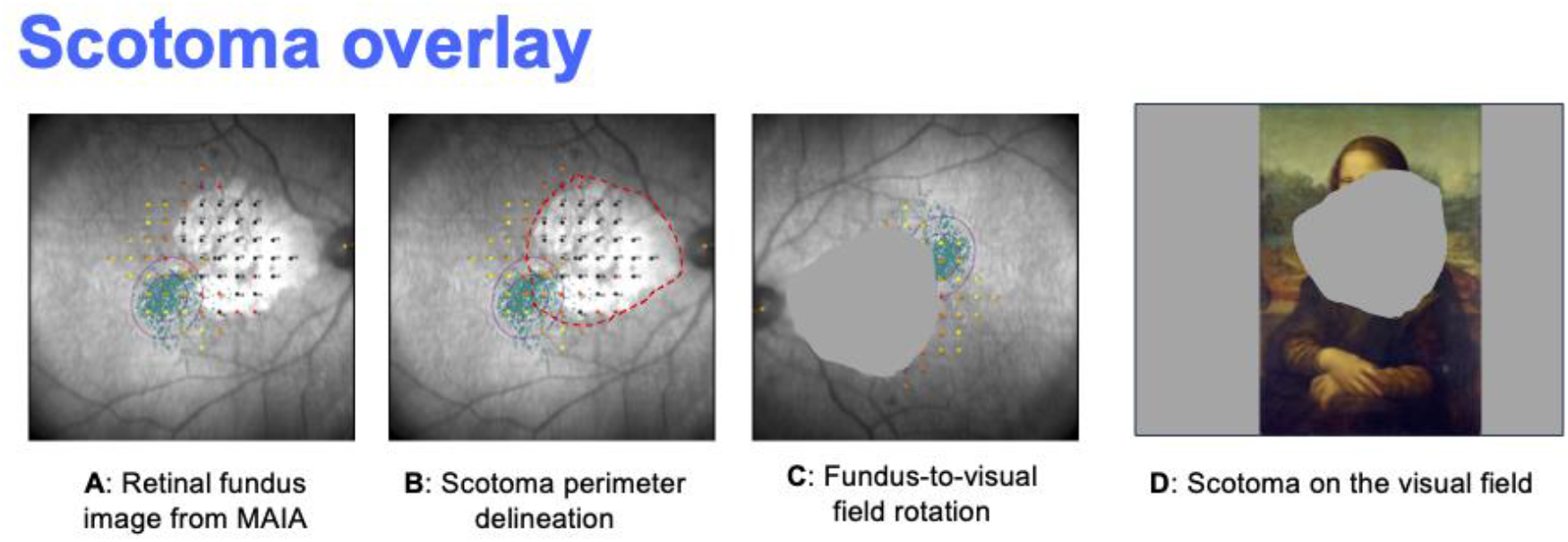
procedure for definition of scotoma overlay on the visual field performed individually for all the MD participants. Macular Integrity Assessment (MAIA; CenterVue, Padova, Italy) microperimetry was used to capture images of the patients’ retinal fundus while they were engaged in a fixation task (**A**). This allowed us to estimate the structural and functional scotoma on the retina and draw its perimeter (**B**). The image was then transformed and rotated to mirror the relationship between retinal grid and visual field (**C**) and used as a gaze-contingent, overlaid scotoma on a computer screen (**D**). Only the preferred eye was included in this analysis, and the non-preferred eye was patched during testing and training. The scotoma border was individually defined for each participant using results from the MAIA exam (Figure 3). The border was determined through a consensus among all authors and based on both anatomical and functional output from the exam.

**Figure 3.**
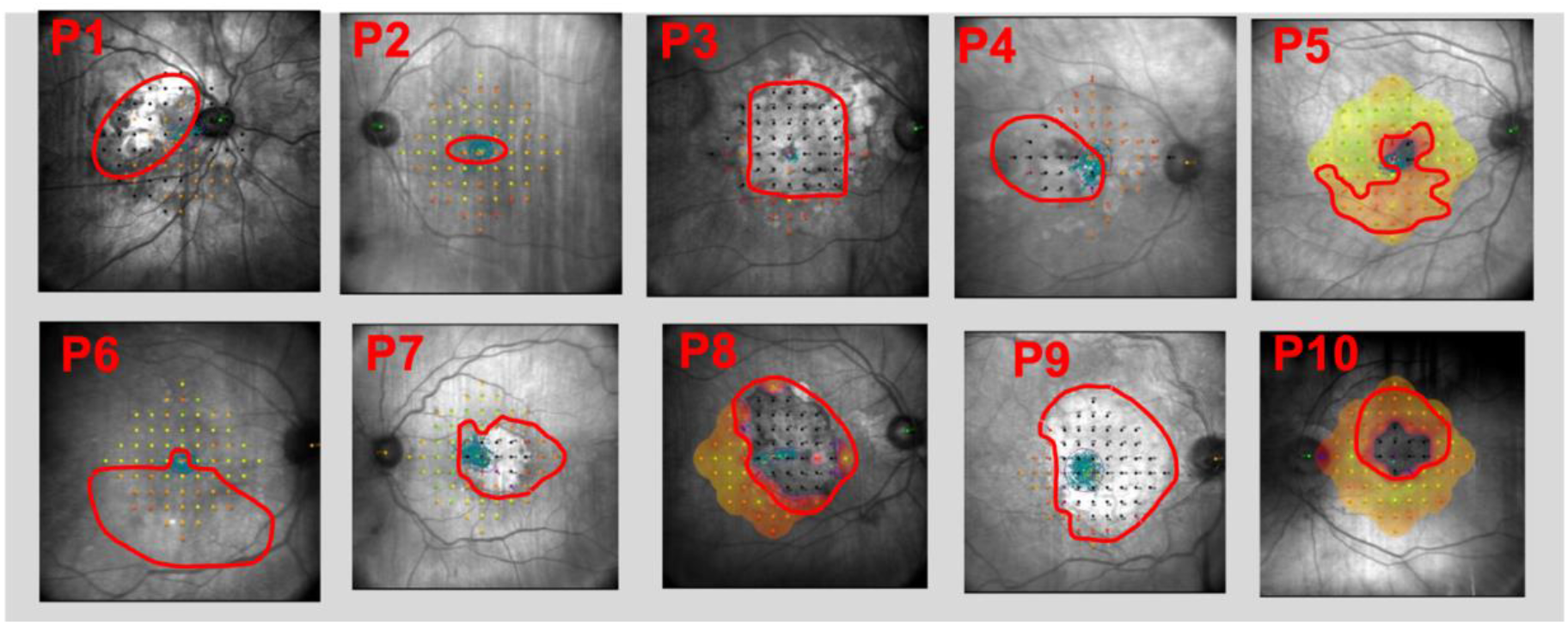
Fundus image from MAIA exam with layout of the scotoma area used for the analysis and the gaze-contingent version of the task.

*Saccadic re-referencing* addresses whether the foveal reference is still present by calculating the percentage of first fixations per trial falling outside the scotoma (i.e., the percentage of first fixations putting the target in a visible location). A value of 100% indicates that no first eye movement placed the stimulus within the scotoma, while 0% indicates that the first fixation in each trial would systematically place the scotoma on the target.

*First saccade landing dispersion* determines whether participants tend to use a consistent location across trials for landing their first saccade after a visual stimulus is presented. The area of first saccade landing locations is computed using BCEA fit encompassing 68% of the total eye-tracking data points, similar to previous studies^14,16,17^.

*Latency of target acquisition* characterized the time interval between the target appearance and the first fixation outside the scotoma.

*Percentage of trials that are useful*, is the percentage of trials in which at least one saccade landed outside the scotoma (“useful” trials, to indicate trials in which the target was visible, thus useful towards performing the task).

*Saccadic precision* addresses the consistency of PRLs across trials by calculating the distribution of locations of the trial’s first fixation that lands outside the scotoma. This fixation could therefore be the first fixation, the second, or the third, etc., and represents the first fixation during which the target can be seen. This visually distinguishes “absolute” first fixations (i.e., first fixations outside the scotoma that happen to be the first fixation in the trial) with a different color from other fixations following initial fixations to the scotoma (see fourth row in Figure 1).

*Fixation stability*, is a within-trial measure of oculomotor stability, characterizing the dispersion of eye positions within a trial.

Importantly, in the case of patients exhibiting multiple PRLs, as revealed by multiple clusters in the saccadic precision analysis (following^14^), these metrics can be extracted separately for subsets of trials in which each PRL is used for the target. In a series of recent papers ^15,18,19^, we used these metrics to evaluate compensatory oculomotor behaviors in a group of healthy individuals trained with simulated central vision loss via gaze-contingent scotoma, showing how these metrics can reveal different eye movement strategies when participants are engaged in a visual task. Additionally, we have demonstrated how these metrics capture changes in oculomotor behavior resulting from visual training^18^.

In the current study, we explore the feasibility of using this oculomotor analysis in MD by examining these metrics in ten patients with central vision loss that were longitudinally studied as part of a vision training intervention. Results revealed diverse and complex patterns of oculomotor behavior, demonstrating the ability of these metrics to capture variability of eye movement behaviors, in patients with central vision loss, thus expanding their use from characterizing oculomotor strategies in simulated scotomata to physiological central vision loss. This suggests that these metrics can improve characterization of oculomotor strategies in patients by separating eye movement behavior into multiple components. This supports our long-term goal to use these metrics to better understand, and ultimately improve visual performance in everyday visual tasks. We discuss opportunities for these metrics to be used to quantify longitudinal changes in oculomotor strategies in patients with MD and how these metrics could provide a sensitive tool to help distinguish between optimal and suboptimal spontaneous adaptations and for evaluating the effects of interventions on peripheral looking strategies.

## Method

### Participants

Ten patients with MD (see Table 1 for demographic info) took part in this study. Patients were recruited through the University of Alabama, Birmingham and UAB Callahan Eye Hospital. Individual characteristics of participants are summarized in Table 1. Inclusion criteria for MD patients were: (1) diagnosis of late-stage MD, (2) Presence of bilateral scotoma near the fovea (3) Severely impaired vision in both eyes (0.7 logMAR or worse), (4) Medical record review indicating this level of disease severity has been present for at least 2 years. These inclusion criteria were chosen to maximize peripheral vision use by ensuring that patients would have considerable central vision loss and rely on peripheral vision for everyday activities. Exclusion criteria are (1) comorbidity with cognitive pathologies, (2) bilateral retinal scotomas larger than 20° diameter. An additional patient with MD dropped at the beginning of the study.

**Table 1:** summary table of MD patients that took part in the study.

| Participant | Age (years) | Tested eye | Visual acuity (logMAR) | Contrast sensitivity (MARS) |
| --- | --- | --- | --- | --- |
| P1 | 72 | Right | .54 | 1.08 |
| P2 | 89 | Left | .92 | .96 |
| P3 | 81 | Left | .58 | .36 |
| P4 | 85 | Right | .86 | 1.24 |
| P5 | 76 | Left | .24 | 1.52 |
| P6 | 86 | Right | .60 | 1.28 |
| P7 | 87 | Right | .52 | 1.44 |
| P8 | 83 | Left | .36 | 1.32 |
| P9 | 87 | Right | .76 | 1.28 |
| P10 | 75 | Right | .94 | 1.16 |

### Stimuli and apparatus

The stimuli and apparatus used included MAIA microperimetry and visual charts for the eye exam (see Procedure below), and computer stimuli and video-based eye tracking for the lab visit (see Procedure below). During the lab visit, participants underwent a visual task described below. During the task, eye movements were tracked at 120Hz with infrared, video-based eye-tracker (TRACKPixx3, VPixx Technologies Inc., Saint-Bruno, Canada). Stimuli, in the form of smiling or frowning emojis, were presented on a Display++ (Cambridge Research Systems, UK). Participants were asked to report whether the emoji was smiling or frowning by pressing the corresponding button on a Vpixx response box. The size of the emoji was set individually for each participant based on pilot testing. Importantly, the primary goal of this task was not to achieve precise matching of differences across participants, but rather to ensure their interaction with a visible stimulus. Testing was conducted monocularly; the untested eye was patched during testing. Viewing distance was 70cm. Participants were tested while wearing their habitual spectacle or contact lens correction appropriate for the viewing distance. As a primary challenge for gaze-contingent displays is achieving the perception of smooth movement of the gaze-contingent component, we evaluated several hardware and software configurations before conducting the study. We then selected the combination that exhibited the shortest latency. Details of this testing are reported in^20^. The tested eye was chosen on a participant basis as the more appropriate for the goal of the assessing PRL and, in future studies, training of promoting PRL development (i.e., dominant eye, large central scotoma, unstable/undefined PRL from the MAIA exam).

### Procedure

Before the lab visit, participants underwent a clinical eye examination at UAB Callahan Eye Hospital. Patients underwent an ocular exam through the Macular integrity assessment (MAIA) microperimeter (CentreVue, Padova, Italy). This microperimetry exam consisted of a fixation cross that the participant is asked to fixate, while targets, in the form of small beams of light, are presented randomly in various locations in the visual field (more details). This exam allows us to evaluate multiple regions in the visual field and generate a visual field integrity map for each participant. Additionally, through this exam, we could estimate fixation stability (defined as both the area of a BCEA fit to a proportion of the overall number of fixations or the % of fixations within a 2° and a 4° radius around fixation target) and PRL location (as the center of the fixation BCEA). Additionally, participants underwent basic visual chart exams, specifically the Early Treatment Diabetic Retinopathy Study (ETDRS) chart (visual acuity) and the Mars Letter Contrast Sensitivity Test (contrast sensitivity). Following the eye exam visit, participants attended a lab visit in which they underwent a 20-minute session of a visual search and acuity task (see below). Before the tasks, a calibration and validation procedure were performed. Specifically, we used a custom MATLAB script adapted from the VPixx calibration script, which incorporated visual aids for fixation, an enlarged initial calibration dot, and slower pacing to better accommodate patients. Visual assessments were conducted before and after each session. A chin-and-headrest was used to ensure that the eyes were aligned with the center of the screen and that the head was stable throughout the session. All the computer-based tests were conducted at a viewing distance of 70cm.

### Visual search and acuity task

Participants were presented with a display showing an emoji, appearing in a random location on a computer screen. Participants were then asked to report whether the emoji was smiling or frowning. The stimulus remained on screen until either the participant provided a response, or the trial timed out (after 8 seconds), whichever occurred first. Throughout the session, a gaze-contingent display, overlaying an opaque form of the shape of each participant’s retinal scotoma was active. In the control condition, participants performed the task without a gaze-contingent scotoma. The stimulus size was kept constant throughout the session and chosen individually to be of a size that was visible for the patient. Since our main focus was to provide an equal amount of eye movement experience in terms of time rather than number of trials and given that some participants found the task challenging even when stimulus size was individually adapted to optimize performance, the number of trials varied across participants in order to keep the overall session duration within 20 minutes. Acoustic feedback was provided to inform participants whether their response was correct, incorrect, or whether the trial timed out.

### Oculomotor metrics

Details concerning the metrics, both agnostic and PRL-specific, are described in^15^. Eye-tracking data was collected at the rate of the gaze-contingent display (120 Hz). Saccades were determined based on eye velocity calculated on a frame-by-frame basis. The velocity threshold for identifying a saccade was 60 deg/s. The microsaccade threshold, used to determine periods of eye stability contributing to fixation, was 20 deg/s. Additionally, a further criterion based on eye position was applied: if eye location changed by more than 1 dva between consecutive frames, this was counted as a breach of fixation even if the velocity threshold was not reached. Fixations were defined as consecutive samples below the velocity threshold for at least 80 ms.

As part of a clinical trial, participants with MD were randomized to two variations of a visual task (a ‘scotoma awareness’ group, which had a visible, gaze-contingent overlay, and a control group, which had no visual constraints. Specifics are discussed in the section ‘*Oculomotor metrics’ ability to capture longitudinal changes in eye movement behaviors (training effects)’* in the result section). Given that the trial is on-going at the time of writing this manuscript, we present data on the entire sample, independent of this experimental manipulation. Details of the training procedures and their differential effects on oculomotor metrics will be reported in a follow-up paper after the clinical trial and unblinding is complete. Given the small number of participants and the large variability both in scotoma characteristics and metric scores, we decided not to perform statistical comparisons between groups. For completeness, we list here the participants who were included in each group: Scotoma awareness group: P1, P2, P3, P5, P8, P10. ‘Invisible scotoma’ (control group): P4, P6, P7, P9.

In the context of the present study, the scotoma overlay procedure served the main purpose of defining the inside vs outside-of-the-scotoma oculomotor behavior used to calculate our oculomotor metrics (See Figure 1). A second use of the scotoma overlay, which will be the focus of a future publication, is to define the shape, size and location of the gaze-contingent component of an oculomotor training procedure we developed (‘scotoma awareness’). While we do present training data here for both the experimental (‘scotoma awareness’) and the control groups (‘invisible scotoma’), our main aim with this paper is to describe the use of new oculomotor metrics to describe eye movement behavior in patients with MD independently from the visual task or number of training sessions they performed. Of note, P1 and P2 took part in an earlier version of the study and underwent a different procedure of scotoma definition that simplified the borders to approximate an ellipse. From P3 onward, we refined the procedure to better capture the heterogeneous shapes of each patient’s scotoma.

## Results

As a first analysis, we extracted the six oculomotor metrics for the whole sample of participants. A breakdown of each metric for each participant, alongside the ‘standard’ fixation analysis, is provided in Figure 4. In rows 1-4, data is presented as fixations in the visual field space, with the scotoma layout projected onto each patient’s visual field.

**Figure 4.**
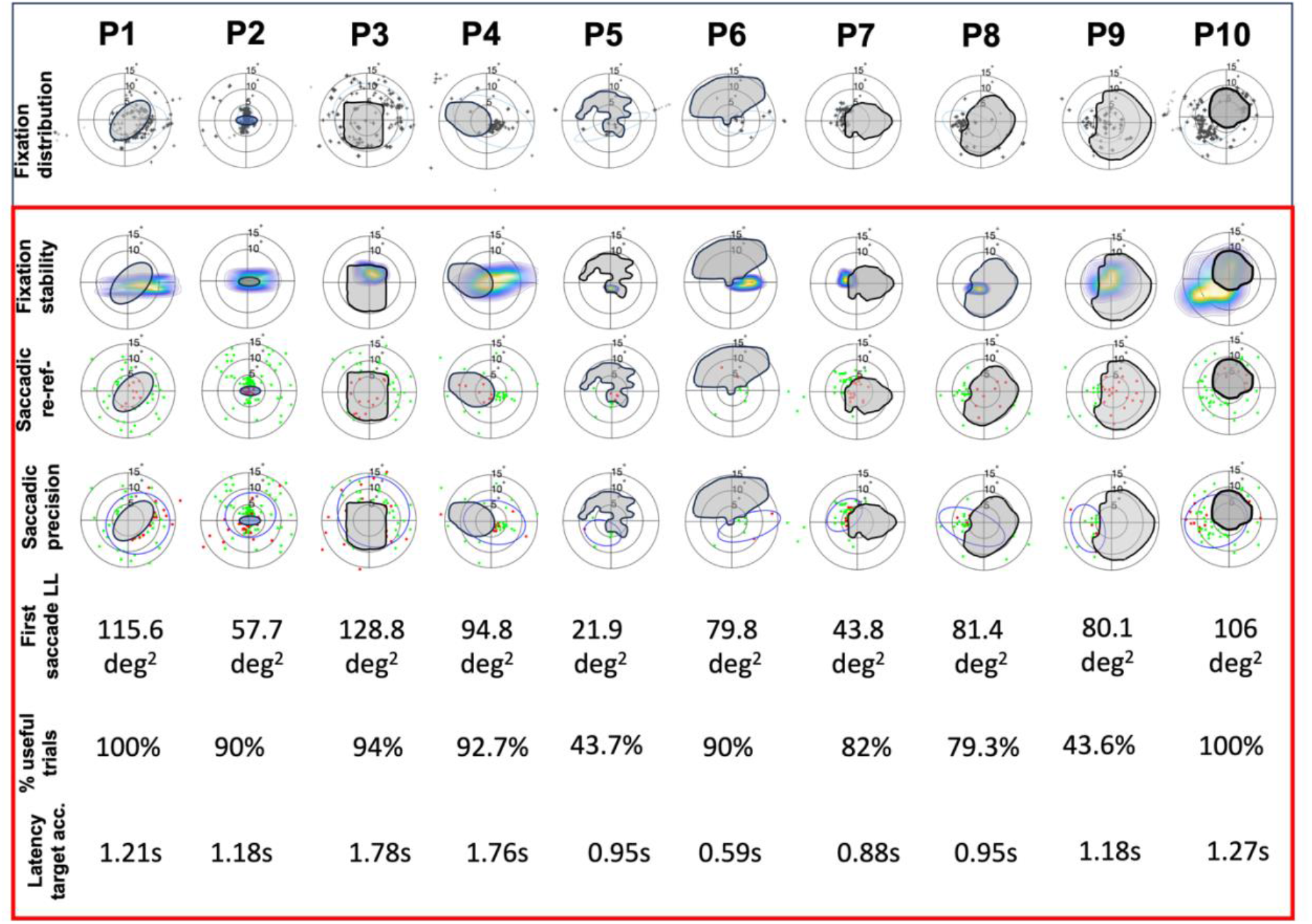
Oculomotor metrics extracted individually for each participant during their first session. In the first row we show a ‘standard’ analysis of fixation distribution as reference. In each graph, the individual scotoma is overlayed on the visual field representation.

To visualize the group data distribution across metrics, Figure 5 presents group-level data for each metric, with individual data points labeled for each participant.

**Figure 5.**
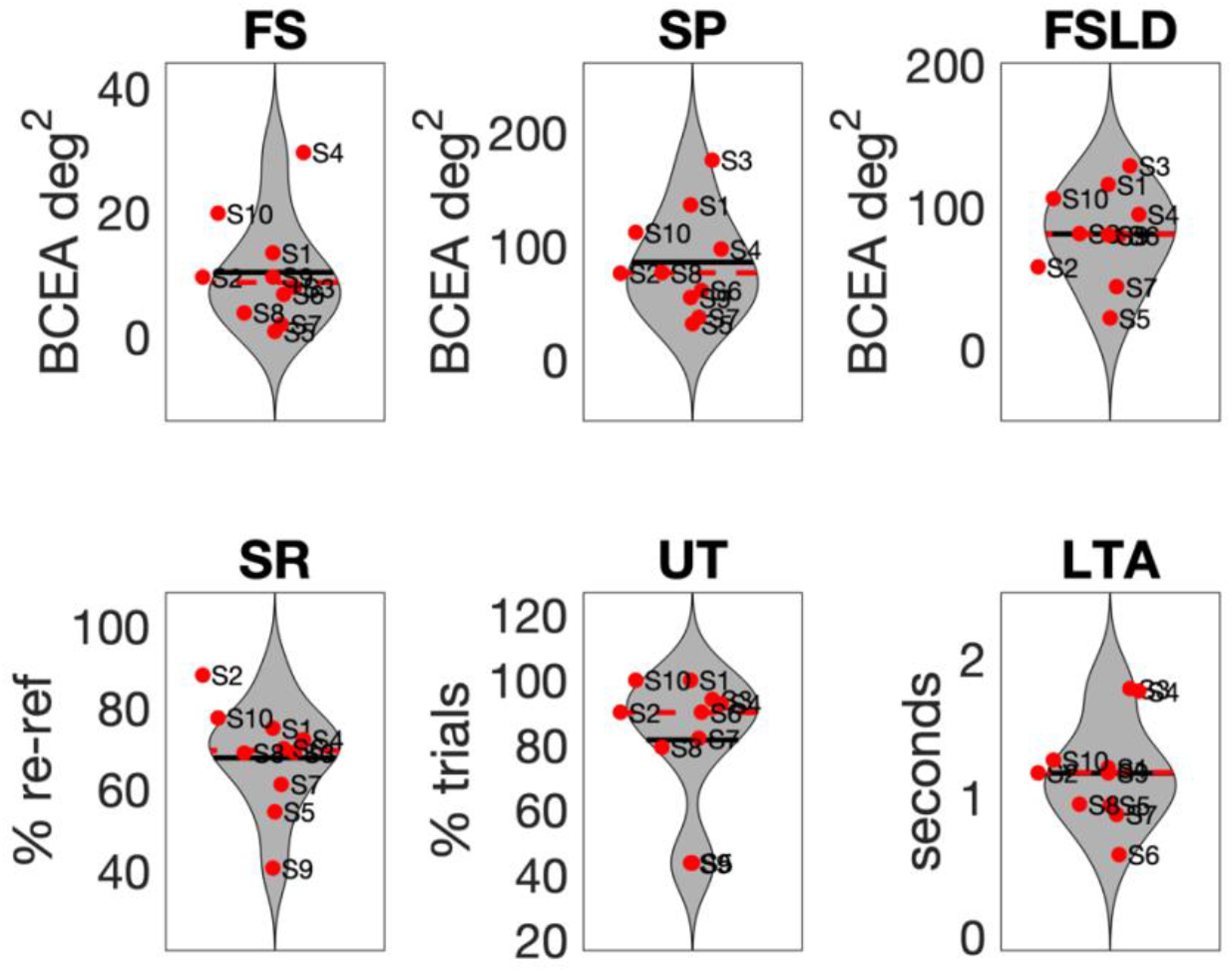
metric distribution across patients. In each graph the black line indicates the mean, and the dotted red line shows the median. (FS: fixation stability, SP: saccadic precision, FSLD: first saccade landing dispersion, SR: saccadic re-referencing, UT: useful trials, LTA: latency of target acquisition).

### Knowledge gained relative to standard fixation analysis

To verify that the oculomotor metrics provide non-redundant information, we assessed whether they capture different aspects of oculomotor behavior through principal component analysis (PCA). Data from the six metrics, measured for each participant (Figure 4), were included in the PCA. These components are shown in Table 2 and explain 61.7%, 19.7%, and 11.3% of the variance, respectively. All metrics have relatively strong weights (> 0.45) on at least one component, indicating that each metric contributes meaningfully to describing participants’ behaviors. Additionally, no single metric dominated any component, suggesting that using a combination of metrics provides more nuanced and non-redundant insights into oculomotor behavior.

**Table 2:** PCA coefficients of components 1, 2, and 3 for the six metrics.

| Metric | Component 1 | Component 2 | Component 3 |
| --- | --- | --- | --- |
| Re-referencing | 0.34 | 0.64 | 0.18 |
| Latency | 0.38 | -0.51 | 0.18 |
| First saccade | 0.45 | -0.2 | -0.39 |
| Saccadic precision | 0.47 | -0.14 | -0.46 |
| Useful trials | 0.43 | 0.48 | -0.03 |
| Fixation stability | 0.37 | -0.2 | 0.75 |

### Dissociation of oculomotor metrics

By breaking eye movement behavior into multiple metrics, we can approximate a molecular approach to eye movement analysis, revealing hidden complexities. A PCA revealed that no two metrics are tightly related, suggesting that our choice of analysis does encompass different aspects of eye movement behavior. This is further confirmed by qualitative observations concerning the metric distributions. For example, while exhibiting similar first saccade landing dispersion (79.9 vs 80 deg^2^), P6 and P9 show different latency, with the former being almost twice as fast at latency of target acquisition (0.6s vs 1.18s). Similarly, while P6 and P3 exhibited comparable saccadic re-referencing (70% vs 72%), the latter was almost three times slower in acquiring the target (0.6s vs 1.78s).

Saccadic precision may be an indicator of a PRL in progress: it is known, both in healthy individuals trained with artificial scotoma and in patients, that oculomotor re-referencing follows the development of a PRL, meaning that patients might use a PRL without necessarily having their oculomotor system shifted toward the PRL (yet). We here observe behaviors in which some participants, such as P1, P4 and P10, exhibit high % of re-referencing (indicating a shift away from the foveal reference) but a large dispersion of saccadic precision (suggesting that the shift away from the fovea has not been accompanied yet by the selection of a specific eccentric location). Conversely, P5 and P9 who show low re-referencing, indicating a persistent foveal referencing, but low dispersion in the saccadic precision metric, which suggests that a PRL might be in development. Finally, there are patients, such as P2, P7, P8 and P9, which show high referencing and low dispersion, indicating the most promising profile (suggesting that the first eye movement after target acquisition places the target away from the scotoma and within a relatively small eccentric region).

### PRL-specific analysis and Use of multiple PRLs in MD

Clinical evidence and reports from clinicians and occupational therapists suggest that MD patients might exhibit multiple PRLs ^2,9–13^. Consequently, an approach that treats fixation distribution unimodally might hide the multimodal nature of the dataset, thus leading to inaccurate estimation of fixational characteristics and PRL locations ^14^.

In a further step in our analysis, we utilize a k-mean approach to distinguish cluster of fixations indicative of the use of multiple PRLs. By doing so, we can better observe compensatory behaviors that might be masked by the conflation with trials without useful fixations. Specifically, we conduct the k-mean analysis on the saccadic precision distribution, thus considering only trials in which at least one ‘useful’ fixation was observed outside of the scotoma. Additionally, for participants exhibiting multiple PRLs, this breakdown can help distinguish whether there exists a “primary” PRL and one or more “secondary” ones. Of note, in the PRL-specific analysis we do not report the metric corresponding to useful trials, as by definition all the trials used in this analysis had at least one fixation outside the scotoma, thus resulting in a score of 100% for all the participants.

While most of the participants showed an overall ‘main’ PRL, our PRL-specific analysis showed that most participants exhibited more than one PRL (Figure 6). This could be due, at least in part, to the randomized nature of the stimulus location. Indeed, patients might use a secondary PRL if the stimulus appears closer to that with respect to their main PRL. Additionally, the breakdown in Figure 6 reveals that most participants showed many ‘useful’ trials (orange + gray areas), suggesting that their spontaneous strategies are overall functional.

**Figure 6.**
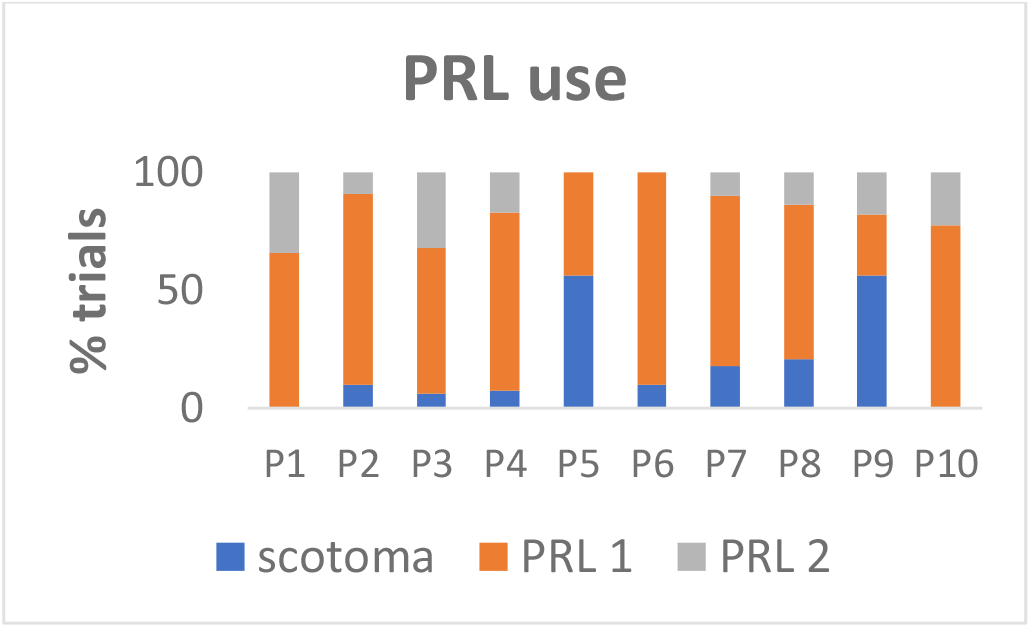
PRL use across trials (from saccadic precision). This graph shows the percentage of trials in which the patient had a first saccade within one (or more) PRL(s) vs trials in which they never made a saccade outside of the scotoma (non-‘useful’ trials). Distinction between patients with one (i.e., P6 and P7) vs two PRLs came from the KDE analysis on the saccadic precision metric (see red box in Figure 7).

To further assess PRL-specific oculomotor behavior across participants, in Figure 7 we present the metric extracted individually for each patient. As in Figure 4, Fixation stability, saccadic re-referencing and Saccadic precision are visualized in the visual field space, with the scotoma layout projected onto each patient’s visual field.

**Figure 7.**
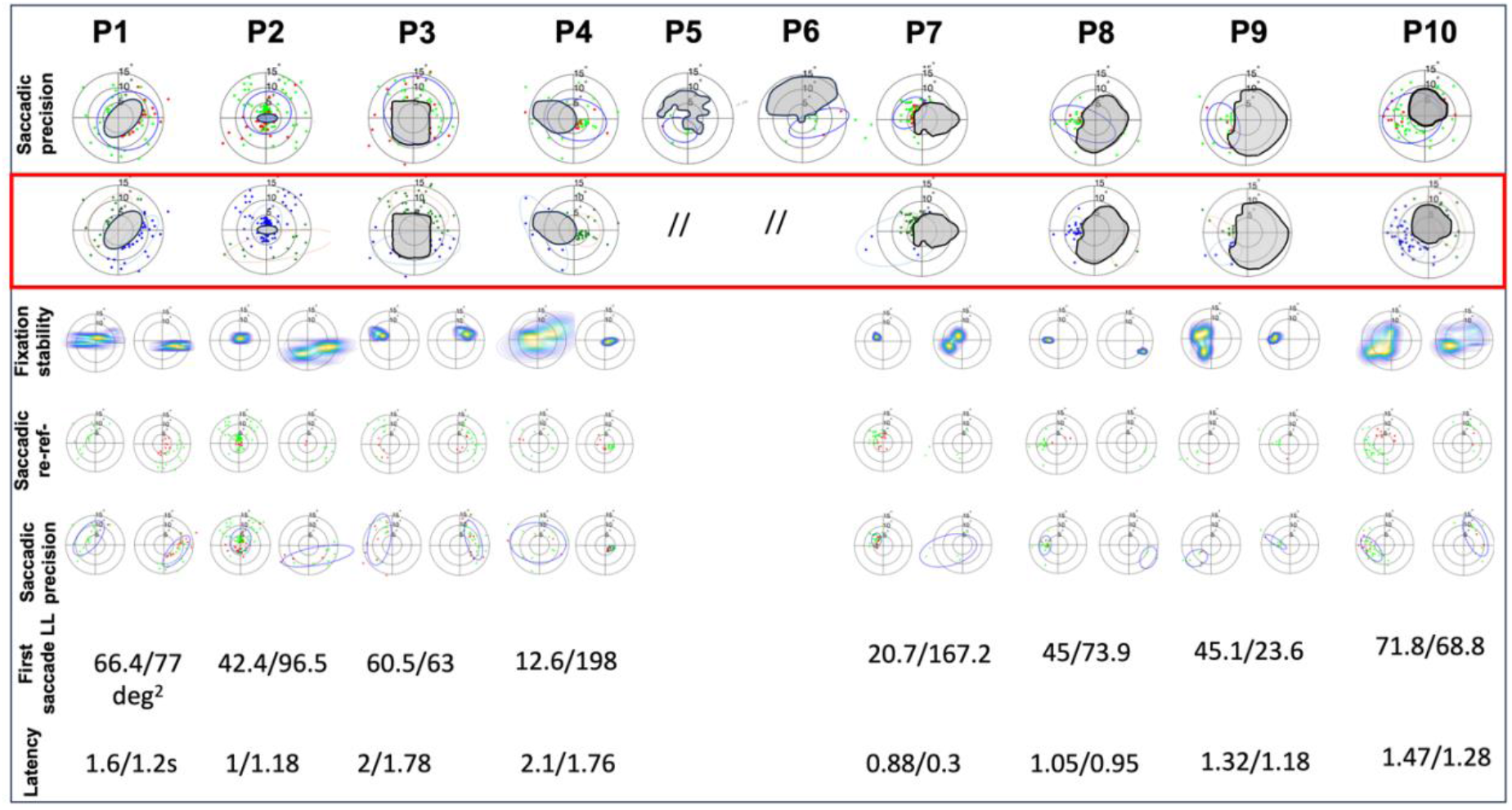
Individual participant data for the oculomotor metrics in the PRL-specific analysis. The first row shows the saccadic precision analysis, whose data is used to conduct the k-mean analysis to test for the use of multiple PRLs (second row). In each of the subsequent rows, the oculomotor metrics are extracted separately for each PRL. Note that P6 and P7 only had one PRL, so subsequent rows’ information is the same as shown in Figure 4, and so is not presented here.

To visually examine the distribution of the group data across metrics, Figure 8 shows group-level data for each PRL-specific metric, with individual data points labeled for each participant.

**Figure 8.**
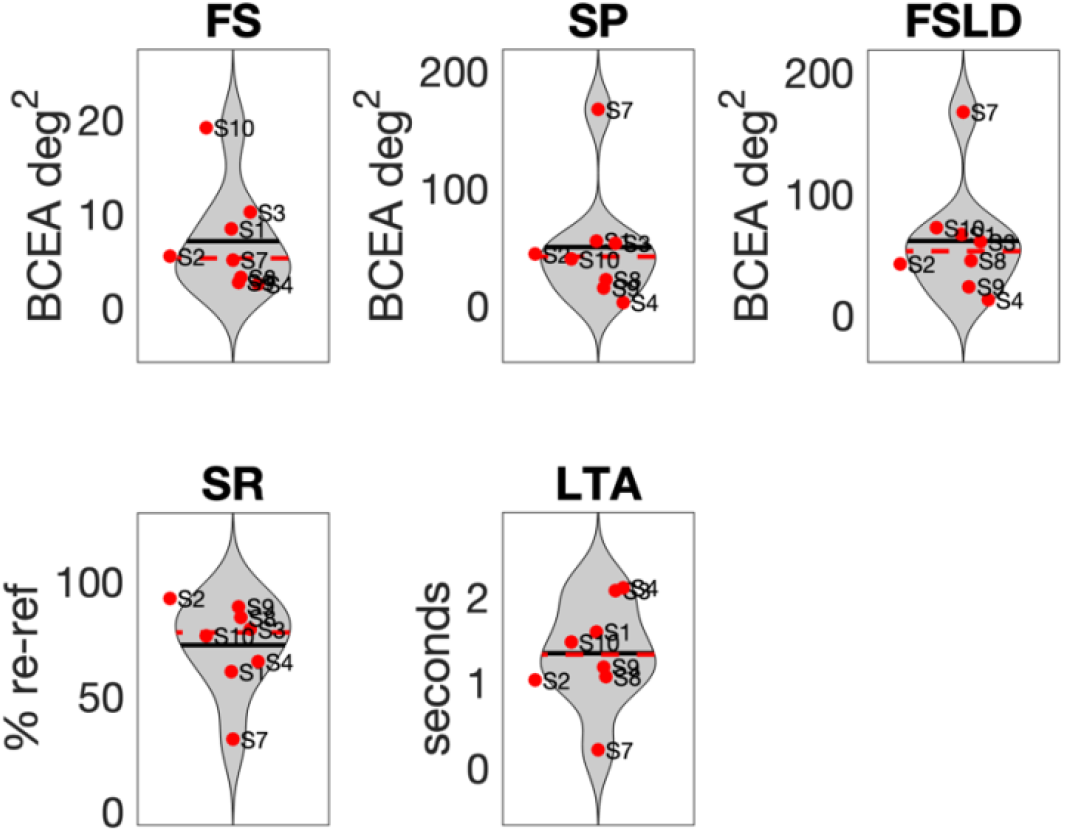
PRL-specific metric distribution across patients. In each graph the black line indicates the mean, and the dotted red line shows the median. (FS: fixation stability, SP: saccadic precision, FSLD: first saccade landing dispersion, SR: saccadic re-referencing, UT: useful trials, LTA: latency of target acquisition).

### Knowledge gain with respect to PRL-agnostic analysis

Previous literature and clinical practice suggest that patients may use different PRLs depending on the characteristics of the stimulus or task ^11,21,22^. Consequently, any analysis assuming a unimodal distribution of eye positions across trials risks conflating distinct strategies associated with different retinal locations. This could result in incorrect estimates of key measurements such as PRL location or fixation stability. To address this possibility, we envisioned an additional step to our analysis: a k-means clustering technique to identify the potential presence of multiple PRLs. Once fixations were classified into different PRL groups, we applied the same metrics separately to each subgroup of trials corresponding to each PRL.

### Oculomotor metrics and scotoma size

As peripheral fixation tends to me less stable than foveal fixation ^23^, one might expect that the size of the scotoma is inversely related to fixation stability. To test whether any of our metrics were affected by scotoma size, we conducted correlation analysis across metrics (see Figure 10). Interestingly, in our sample of participants, most metrics do not seem related to scotoma size, suggesting that oculomotor adaptation to central vision loss might be multidimensional and complex process, an analysis needs to take into account multiple factors. Nonetheless, we found a significant correlation between saccadic re-referencing and scotoma size, with participants with smaller scotomas showing larger % of trials with a first saccade outside of the scotoma.

**Figure 10.**
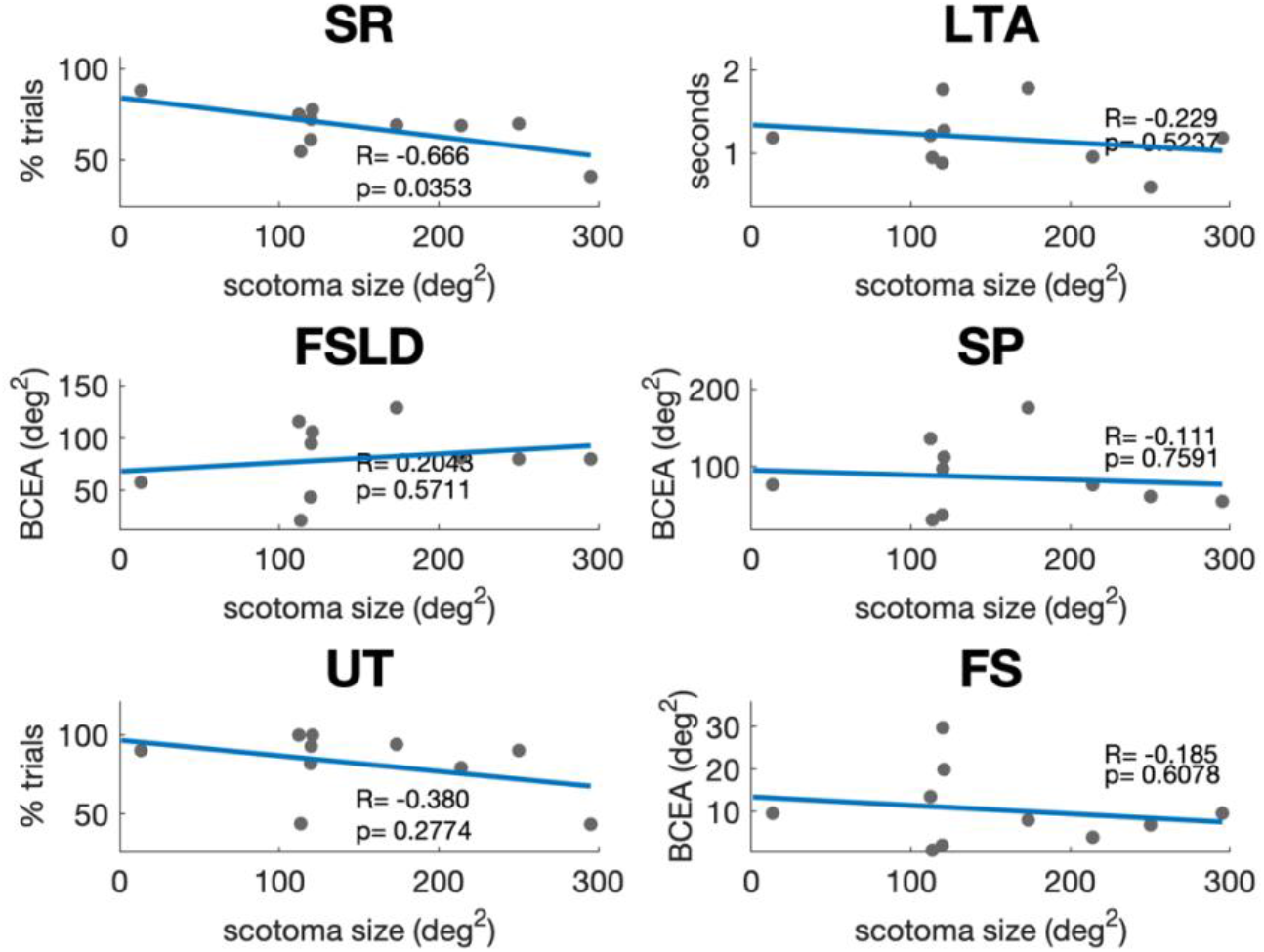
Correlation between scotoma area and oculomotor metrics across participants.SR: saccadic re-referencing; LTA: latency of target acquisition, FSLD: first saccade landing dispersion; SP: saccadic precision; UT: percentage of useful trials; FS: fixation stability.

### Comparison with simulated scotoma

Simulating central vision loss in healthy individuals through the use of eye tracker-guided, gaze-contingent display has been recently used as a framework for the study of compensatory oculomotor strategies in central vision loss, as well as a template for testing possible rehabilitative intervention in MD ^17–19,24–29^. One of the main goals of this paper was to extend the application of these oculomotor metrics, initially developed in healthy individuals using simulated scotomas, to patients with central vision loss. Comparing metrics distribution in simulated (from Maniglia et al., 2020) and physiological scotomas, as in Figure 11, we observe some interesting patterns: patients with MD outperform healthy individuals trained with simulated scotoma in re-referencing, both at baseline (independent t-test between MD and healthy individuals scores: t_27_= 7.09, p < 0.001) and after training (independent t-test: t_27_= 2.8, p = 0.01). Somehow surprisingly, patients show a larger first saccade landing location (vs pre, independent t-test: t_27_= 7.36, p < 0.001, vs post, independent t-test: t_27_= 6.63, p < 0.001), suggesting that they were not systematically using a PRL across trials. This is consistent with the idea that the PRL that emerges from static fixation tasks, such as the microperimetry exam used to characterize each patient’s PRL at the onset of the study, might not reflect eccentric fixation behaviors in re-fixational tasks (Chung, 2011). In terms of percentage of useful trials, patients have significantly higher useful trials when compared to simulated scotoma before (independent t-test: t_27_= 2.39, p = 0.024) but not after training (independent t-test: t_27_= 1.29, p = 0.2), suggesting that training in healthy individuals improved useful trials. Interestingly, while at baseline patients and healthy individuals trained with simulated scotoma exhibited similar latency to target acquisition (independent t-test: t_27_= 0.98, p = 0.334), the latter outperformed the former after training (independent t-test: t27= 2.93, p = 0.007). Similarly, after training, but not before, controls had higher saccadic precision (vs pre, independent t-test: t_27_= 1.74, p = 0.93, vs post, independent t-test: t_27_= 3.77, p < 0.001).

**Figure 11.**
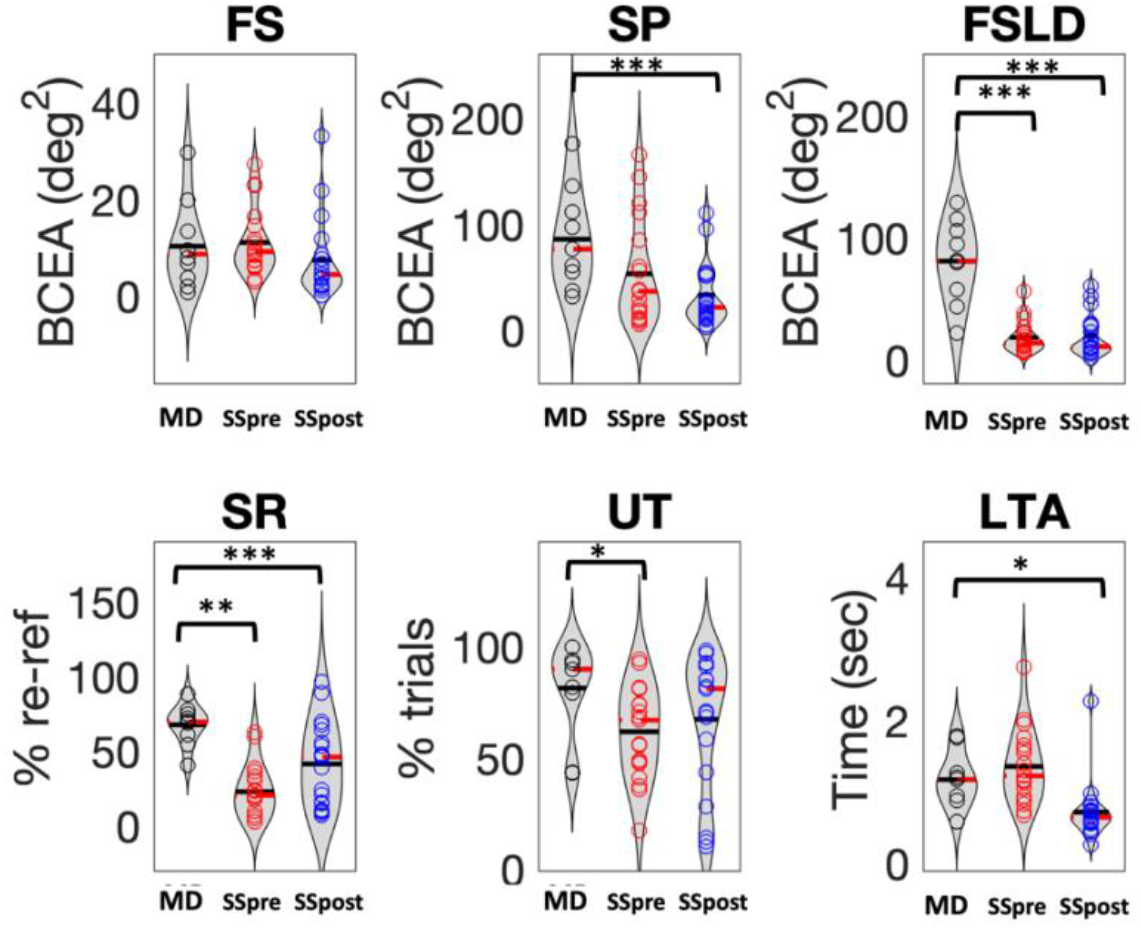
comparison of oculomotor metrics between patients and healthy individuals tested on a simulated scotoma display (data from Maniglia, Visscher and Seitz, 2020). In each graph the black line indicates the mean, and the dotted red line shows the median. (. (MD: patients with MD, SSpre and SSpost: simulated scotoma participants before and after training, respectively; FS: fixation stability, SP: saccadic precision, FSLD: first saccade landing dispersion, SR: saccadic re-referencing, UT: useful trials, LTA: latency of target acquisition).

### Oculomotor metrics’ ability to capture longitudinal changes in eye movement behaviors (training effects)

A key aspect of these metrics is their potential for evaluating training interventions. While most training studies in patients with MD focus solely on behavioral outcomes or simple fixation analysis, these metrics enable a multidimensional examination of eye movements. To illustrate their ability to capture training-related changes, Figure 12 presents the output of two metrics— Fixation Stability and Saccadic Re-referencing—at the beginning, midpoint, and end of a training protocol aimed at improving patients’ awareness of their scotoma border. In addition to their respective scores, a visual inspection of these metrics provides a clearer understanding of the spatial distribution of eye movements and the use of peripheral vision across participants and training sessions. Specifically, some participants (e.g., P3, P4, P9, P10) maintained a broad use of their peripheral vision, while others (e.g., P5, P6, P7) exhibited shifts in its location. Furthermore, applying a KDE analysis to the fixation stability metric helps determine whether participants exhibit a single fixation stability cluster or multiple ones.

**Figure 12.**
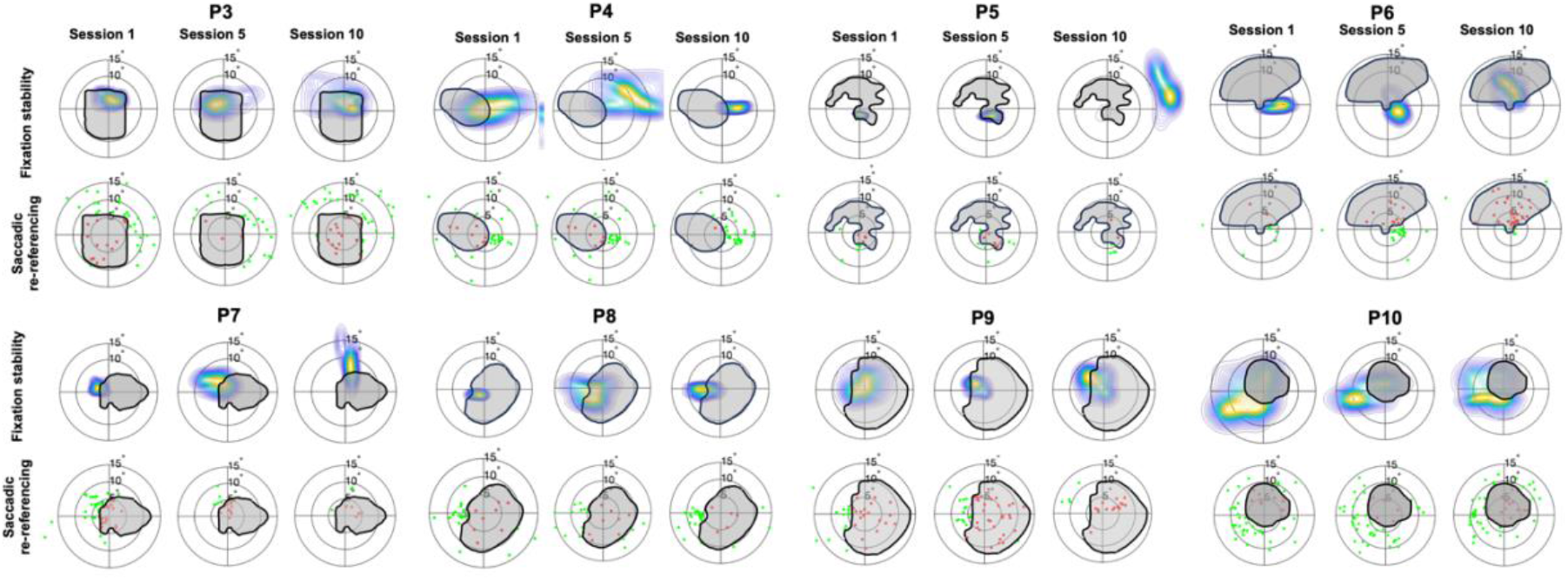
Saccadic re-referencing and fixation stability at the beginning, halfway through ad on the last session of the training.

## Discussion

Macular Degeneration (MD) represents a serious health concern worldwide, and the main cause of visual impairment in the western world ^1^. Late-stage MD leads to loss of central vision, forcing patients to use their peripheral vision to perform everyday tasks such as navigation, reading, face recognition and more ^3,8,30^. A large majority ^6,7^ ) of these patients rely on a specific region in the spared peripheral vision to perform such tasks, the Preferred Retinal Locus (PRL) which they use as a surrogate fovea ^4,5,31^. Characterizing the way patients with MD use their residual vision can help better understand spontaneous mechanisms of oculomotor compensation for central vision loss, distinguish between advantageous and suboptimal spontaneous strategies, and evaluate rehabilitative training interventions. However, current standard assessments of oculomotor behaviors seem to offer a simplified, and possibly misleading, evaluation of PRL location and characteristics, due to their reliance on basic fixation tasks that might not represent how patients use their peripheral vision across different tasks, stimuli, and environmental conditions. For example, a standard way of defining the PRL is the center of a BCEA fitted on a certain percentage of the overall fixation distribution, but evidence of multiple PRLs would mean that such a unimodal approach to the fixation distribution might be misleading. Indeed, this issue has been observed before ^14^.

As a first step towards a more complete characterization of oculomotor behavior in conditions of simulated central vision loss, in 2020 we introduced six metrics with which evaluate eye movement strategies while participants are engaged in a visual task ^15^, and we extracted those metrics in a group of healthy individuals trained with simulated scotomas ^18^. In this study, we present data from 10 patients with MD as proof of concept and evidence of feasibility the use of these metrics to characterize different profiles of eye movements in central vision loss. The goal of these metrics is to be able to capture the heterogeneity of the visual experience in MD, which is characterized by vast differences in size, location and shape of the scotoma, and onset and comorbidity. Specifically, our analysis addresses three major issues associated with standard fixation analysis approaches.

First, standard fixation analysis typically relies on a single measure — most often fixation dispersion, such as the bivariate contour ellipse area (BCEA) calculated on a certain proportion of the fixation data. This single-dimensional approach may yield similar results across patients with distinctly different eye movement patterns, oversimplifying the complexity of their strategies. Our analysis expands the assessment of oculomotor behavior to six independent metrics, confirmed by principal component analysis to capture distinct aspects of eye movement behavior.

A second issue with standard fixation analysis is that it conflates eye movements over time, thus losing the temporal dimension of oculomotor behavior. While recent studies have begun to address temporal components of simple fixation task ^32^, a systematic temporal analysis, especially in the context of a more dynamic tasks than fixation, has not been widely adopted yet. To address this issue, we developed metrics characterizing temporal aspects of oculomotor behavior, such as dispersion of the first saccade and fixation stability after target acquisition. This breakdown uncovers behaviors that might be overlooked by a general fixation analysis. For example, examining Participant 3’s fixation analysis in Figure 4, which shows a wide distribution, might suggest that this participant has poor fixation stability. However, our fixation stability metric reveals that once the target is acquired, this participant maintains a relatively steady fixation, thus suggesting that the impression of poor stability might depend on a searching behavior once the target appears on screen.

A third issue with the standard fixation analysis concerns the unimodal assumption of eye movement distributions. Most fixation exams, such as those conducted with microperimetry devices (i.e., MAIA, Nidek MP-1), assume unimodal distribution of eye movements. However, this approach may conflate different eye movement strategies, such as the use of multiple PRLs (e.g., Crossland et al. (2005). For instance, a patient systematically using one PRL to the left and one to the right of a central scotoma, would have their BCEA inaccurately centered in between the two true PRLs, incorrectly suggesting persistent foveal referencing, when in fact, most fixations occurred outside the scotoma. This can be seen in the first row of Figure 4, especially with Participants 3 (P3) and P10. For P3, a BCEA analysis would incorrectly place the PRL near the fovea, despite most fixations being outside the scotoma. For P10, the BCEA center would be closer to the scotoma’s edge than to the true, denser cluster of fixations. A kernel density estimate (KDE) analysis, as recommended by ^14^ and adopted in our study, addresses this by revealing complex behaviors that a unimodal analysis would overlook. We further address this issue by adding an additional step of analysis, described as ‘PRL-specific’.

Importantly, we present longitudinal examination of eye movement behavior across these metrics. Participants in this study underwent 10 training sessions, and metrics were extracted for each participant and each training session. Results show a diversity of behaviors across participants that standard fixation analysis might conflate. Future studies in the works will look at the relationship between these metrics, characteristics of the training protocol, and behavioral performance, with the goal of further understanding whether specific oculomotor strategies present themselves as more promising training protocols.

In regard to the training condition of the current trial, the core idea behind our training procedure (“scotoma awareness”) is to leverage evidence from the simulated scotoma framework, which shows that sharp-bordered gaze-contingent scotomas are advantageous for promoting the development of PRLs in healthy individuals. In this sense, we acknowledge that the physiological scotoma, by its nature, does not tend to have sharp edges. Our scotoma served two purposes: on one hand, it created the overlaying shape that participants saw in the gaze-contingent display to accelerate the development of their PRL or improve their use of peripheral vision; on the other hand, the scotoma perimeter allowed our metrics based on inside/outside ratios to be calculated. Of note, the clinical trial testing the extent to which scotoma awareness led to be improved training outcomes is still in progress and we plan a subsequent paper detailing the trial once the study is complete and data are fully unblinded.

### Limitations

In our tasks, participants were allowed to wear their habitual correction (spectacles or contact lenses) or not, depending on their comfort with the computer-based task. While this approach likely increased compliance, it may have introduced a potential confound. Specifically, participants who chose not to wear correction may have exhibited longer and less stable fixations, as well as an increased number of saccades, due to reduced image clarity. Nonetheless, we observed substantial inter-individual variability in eye movement behavior that likely exceeds the differences attributable to refractive correction. Future studies might compare within-participant differences in oculomotor behavior between corrected and uncorrected viewing modalities and test whether our metrics are able to capture them. Additionally, the measurements in this study were conducted monocularly, as the microperimetry exam used in clinical practice are typically monocular, and consequently the PRL assessment tends to be performed monocularly. However, there is evidence that patients might adopt different PRLs when changing viewing condition from monocular to binocular^33–35^. Future studies, especially in the context of ecological validity, should consider using binocular gaze-contingent displays.

There are important differences between simulated and physiological central vision loss, for example visible borders, amount of exposure to the simulated display, etc ^36^. These differences may be in part responsible for the different behaviors observed in Figure 11.

A larger dataset is necessary to further test the feasibility of the simulated scotoma as a framework to study MD in lab, clearly distinguish the different peripheral viewing strategies that seem to emerge from this study, and to determine how these strategies are influenced by various training conditions or stimuli. Future studies could also apply these metrics in machine learning algorithms to identify distinct profiles of eye movement compensation.

While longitudinal analysis is here showcases the ability of these metrics to capture changes in oculomotor behavior across multiple dimensions, there remain questions of the extent to which they relate to behavioral improvements or training condition. For example, while the specific outcome of the training (shown in Figure 12) in terms of changing in the metric scores could at least in part be affected by the specific characteristics of the training procedures participants underwent (‘scotoma awareness’ vs ‘invisible scotoma’). We note that the increased regulation of clinical trials provides some challenges to early reports of unblinded data^37^. We hope soon to publish a follow up manuscript, focused on howe these changes might be specifically related to the training conditions.

### Conclusion

Overall, the present results highlight the diverse peripheral viewing strategies used by patients with MD, both within a single session and across training. This further highlights the need for finer analyses to provide a more detailed characterization than standard eye exams typically offer. Moreover, these metrics appear to be relatively independent from the size of the retinal damage, which aligns with our framework that oculomotor compensation varies significantly between individuals ^38^.

This study, alongside recent publications from our lab using the simulated scotoma framework ^18,19,21,26^, highlight the importance of using appropriate analyses and task when aiming at capturing oculomotor behavior in central vision loss. Indeed, previous evidence suggests that both patients with MD and health vision individuals trained with simulated scotoma might use multiple PRLs, and that the choice of PRL at a given moment might depend on characteristics of the stimulus or the task^7,11,21^. We believed that designing appropriate metrics to capture eye movement behavior in low vision goes hand in hand with designing appropriate tasks that promote a variety of oculomotor behavior, ideally closer to the everyday experience of the patients (including visual search, fixation stability, and object recognition). Indeed, a task involving eye movements may approximate the everyday oculomotor experience of a patient more than a fixation task. This effort towards developing more diverse and ecological measures of eye movement strategies may help shed light on the development of the PRL and help characterize whether patients use a single PRL across tasks and stimuli or switch them depending on the context. Additionally, employing a variety of tasks and stimuli can help disentangle the factors contributing to PRL selection^39^.

Support for the use of gaze-contingent display as a proxy for the study of MD comes from evidence that some of the typical compensatory oculomotor behavior exhibited by MD patients can also be observed in healthy individuals trained for several hours with simulated scotoma, in particular the development of a PRL ^17^ and, after further hours of training, peripheral saccadic re-referencing. While several aspects of the simulated scotoma framework differ from the everyday experience of MD patients, some oculomotor behavior typical of MD have been observed in healthy individuals trained with gaze-contingent scotomas, including the development of a PRL ^17,18^. A partial support for the use of simulated scotomas as a framework for the study of compensatory eye movement strategies in central vision loss comes from the PCA result reported above. In line with what observed in healthy individuals tested with simulated central vision loss^18^, these metrics show similar independence and correlation with principal components.

This is, to the best of our knowledge, the first study to compare eye movements behavior between patients with MD and healthy individuals trained with simulated scotoma across a number of oculomotor metrics. Despite larger individual differences in the MD cohort with respect to the gaze-contingent participants in terms of age, duration of the pathology and scotoma size, shape and location, variability within participants is comparable across the two populations. Interestingly, the overall profile of metric distribution in participants trained with simulated scotoma appeared closer to that of patients with MD after training, consistent with the idea that these metrics can be improved with (spontaneous or lab-based) training ^19^.

Interestingly, as shown in Figure 11, fixation stability did not significantly differ between the MD and the control participants, whereas each of the other metrics showed some significant difference between patients and controls. The measures with the largest differences involved saccades to a target: first saccadic landing dispersion, and saccadic precision. Both metrics measure the precision of saccades to an eccentric target, and in both cases the MD participants, who have had years of experience, show more precise performance. This pattern of results is consistent with the idea that fixation stability may be quickly learned by participants in a simulated scotoma paradigm, whereas altering the mechanisms responsible for planning a saccade may require longer and more consistent training than simulated scotoma can provide.

In conclusion, multiple indices of eye movement behavior can, and should, be identified in patients with central vision loss. By independently assessing these different aspects of oculomotor strategies, we can better understand compensatory eye movements in central vision loss. In future studies, we plan to characterize how these metrics are modified by visual training, similar to what our lab did with healthy individuals trained with simulated central vision loss ^15,18,19^. This paper characterizes the patterns of eye movements present in participants with central vision loss. This work, and the use of these eye movement metrics can pave the way to a better understanding of mechanisms of spontaneous compensatory oculomotor behavior. Development of more effective eye movement training for low vision rehabilitation will rely on such fine scale understanding of a patient’s current and optimal oculomotor behavior.

## DECLARATIONS

### Conflicts of interest/Competing interests

no conflict of interest

### Ethics approval

The study was conducted in accordance with the Declaration of Helsinki (1964) and approved by the University of Alabama at Birmingham ethics committee

### Consent to participate

all participants provided a written consent form and were compensated for their participation in the study

### Consent for publication

all participants provided a written consent form

### Availability of data and materials

data is available on the Open Science Framework (OSF) at the link: https://osf.io/zg8xu/?view_only=9bc54f6845c840c9afe60f487c07c30d

### Code availability

custom-made Matlab code will be available upon request

### Funding

This research was supported by the National Eye Institute (NEI) under grant number R21EY033623-01.

## Notes

### Competing Interest Statement

The authors have declared no competing interest.

